# A Stochastic Model for Cancer Metastasis: Branching Stochastic Process with Settlement

**DOI:** 10.1101/294157

**Authors:** Christoph Frei, Thomas Hillen, Adam Rhodes

**Affiliations:** Department of Mathematical and Statistical Sciences University of Alberta, Edmonton, Alberta, T6G2G1, Canada

**Keywords:** branching processes, birth-jump processes, cancer metastasis, integro-differential equations, reproduction number

## Abstract

We introduce a new stochastic model for metastatic growth, which takes the form of a *branching stochastic process with settlement*. The moving particles are interpreted as clusters of cancer cells while stationary particles correspond to micro-tumors and metastases. The analysis of expected particle location, their locational variance, the furthest particle distribution, and the extinction probability leads to a common type of differential equation, namely, a non-local integro-differential equation with distributed delay. We prove global existence and uniqueness results for this type of equation. The solutions’ asymptotic behavior for long time is characterized by an explicit index, a *metastatic reproduction number R*_0_: metastases spread for *R*_0_ > 1 and become extinct for *R*_0_ < 1. Using metastatic data from mouse experiments, we show the suitability of our framework to model metastatic cancer.

## 1. Introduction

Metastasis is the leading cause of cancer related deaths. It is the process by which cancer spreads from a primary site to distant secondary sites. Because of its implication in over 90% of all cancer-related deaths (Gupta and Massague, 2006; Valastyan and Weinberg, 2011), it is recognized as one of the *hallmarks* of cancer (Hanahan and Weinberg, 2011). The *metastatic cascade* (Gupta and Massague, 2006; Valastyan and Weinberg, 2011; Riggi et al., 2018) provides a mechanistic description of the process, viewing it as an ordered sequence of biological events. Growth of the primary tumor leads to local invasion of surrounding normal tissue. Such local invasion eventually encounters a vessel of the lymphatic or circulatory system. Individual cancer cells — or small clusters of cells — can enter these vessels (intravasate), thereby gaining access to rapid transport throughout the body. If these circulating cells survive their journey and exit the vessel at some distant site (extravasate), they may be able to establish a micrometastasis. Evasion of local defenses and adjustment to the hostile foreign environment can see the *micro* metastasis grow into a *macro* metastasis. Further growth may eventually lead to a secondary tumor.

The stochastic nature of metastasis makes experimental studies difficult, meaning that a thorough understanding of the process is lacking. While some of the steps in the metastatic cascade are well understood — for example, growth and invasion of primary tumors are well studied both experimentally and theoretically (see the reviews in Riggi et al. (2018); Scott et al. (2013)) — understanding of others remains elusive. Travel *to*, and establishment *at* secondary sites are particularly poorly understood, with many theoretical investigations ignoring these aspects altogether. Though recent results (Kaplan et al., 2005) have led to novel theories that metastasis may be more intricately orchestrated than previously thought (Shahriyari, 2016; Rhodes and Hillen, 2019), the process is still believed to be largely stochastic.

Based on these observations, we introduce a new stochastic framework for metastatic spread in the form of a *branching stochastic process with settlement*. This model captures simultaneously temporal and locational dynamics. Stationary tumors emit, or shed, small clusters of cells into the vasculature at random times. These shed cells can then move randomly through the body, modeled by a spatial stochastic process. The moving clusters — also known as circulating tumor cells/clusters (CTCs) — can settle randomly according to a given rate. If the CTCs successfully settle, they may establish a secondary tumor, which itself may shed new CTCs into the blood stream. Both moving and stationary groups of cancer cells die, each at their own rate.

In contrast to existing stochastic metastasis models, our framework accounts for both travel between primary and secondary sites and establishment at the secondary sites. Hartung and Gomez (2014) propose a stochastic model with secondary metastatic emission as a cascade of Poisson point processes and link it to the deterministic model introduced by Iwata et al. (2000). Differently from us, Hartung and Gomez (2014) model only the size development over time and not the location.

From a probabilistic viewpoint, our model represents a branching process with general dynamics and including settlement and death of particles. Branching Brownian motion has been analyzed for more than fifty years, starting with seminal work (Ikeda et al., 1968; McKean, 1975) on the fundamental link between branching Brownian motion and the Fisher-Kolmogorov-Petrovsky-Piscounov equation. Since then branching Brownian motion has been intensively studied in its own right (Biggins et al., 1991; Kimmel and Axelrod, 2015). In statistical mechanics, branching Brownian motion is used for models of spin glasses (Bovier, 2016). In biology, branching processes have been applied in a range of areas such as molecular biology, cellular biology, and human evolution (Haccou et al., 2005; Jagers, 1975; Kimmel and Axelrod, 2015). Recently, branching processes have been found useful in simulating semi-linear partial differential equations (Henry-Labordere et al., 2014, 2017). To the best of our knowledge, this paper is the first to introduce branching processes with settlement and study their applications in the context of cancer metastasis.

For the branching stochastic process with settlement, we prove a characterization of the following key quantities: the expected location *F* of metastases, their locational variance *V*, the distribution *H* of the furthest invading metastasis, the metastatic extinction probability *Q*, and the ratio of moving versus stationary cancer cell groups. We show that each of the first four quantities — *F,V,H*, and *Q* — satisfies a non-local integro-differential equation with distributed delay. These non-local differential equations are of the same type for all cases and are non-linear in some cases. Generalizing the renewal equation, integral equations for age-structured branching processes were already derived in Kimmel (1982), where a connection to dynamical systems and control was made. In our paper, the integral equations employ spatial non-local terms and distributed delays, and also have a close connection to dynamical systems. From their derivations, we obtain existence of solutions to the integro-differential equations specific to *F,V,H*, and *Q*, but we go beyond this, and present a detailed analysis of a general type of such non-local integro-differential equations, including positivity, and local and global existence of solutions.

We also provide a simple classification of the asymptotic behavior of branching stochastic processes with settlement. We identify a *metastatic reproduction number R*_0_, which is explicitly given in terms of the rates of metastatic shedding, settlement, and death. For completeness, we present its derivation in two ways: from general results for two-type branching processes and from a stability analysis for differential equations. In particular, we provide a global stability result of extinction of metastases for *R*_0_ < 1 and growth for *R*_0_ > 1. By parametrizing the model on experimental mouse metastatic data, we numerically explore the resulting metastatic pattern and confirm the suitability of our framework to model metastatic cancer.

The remainder of this paper is organized as follows. Section 2 introduces the model, presents the results and relates the stochastic model to integro-differential equations. In Section 3, we analyze this type of integro-differential equations in more detail and characterize their asymptotic behavior using the metastatic reproduction number. Section 4 shows how our framework can be applied in the context metastatic cancer while Section 5 concludes and discusses possible extensions.

## 2. Branching stochastic processes with settlement

We define the *branching stochastic process with settlement* as follows: We start with one tumor that is located at locational position 0. We assume that the tumor sheds individual cells and small groups of cells into the circulatory system. It is believed that such CTCs are most responsible for metastasis formation (Friedl and Mayor, 2017). Hence, in this model, we focus on the shedding of cell clusters. At a random time *ν*, the primary tumor emits a cell cluster, which starts moving randomly. We model the movement of this cell cluster by a *d*-dimensional stochastic process (*B*(*t*))_*t⩾*0_ with cumulative distribution function *G*(*t,x*) = *P*[*B*(*t*) ⩽ *x*] for *x* ∈ ℝ^*d*^ and *t* ⩾ 0. Here and in the following, inequalities between vectors are understood element-wise: *y* ⩽ *x* for *x,y* ∈ ℝ^*d*^ if *y*^*i*^ ⩽ *x*^*i*^ for all *i* = 1, …, *d*. We assume that for every fixed *t, G*(*t*,.) is absolutely continuous so that there is a density function *g*(*t*,.) with 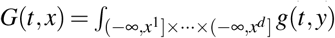 d*y* for all *x* ∈ *ℝ*^*d*^. An example for (*B*(*t*))_*t⩾0*_ is a *d*-dimensional Brownian motion, in which case *g* is the multivariate normal density function 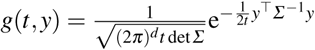 for a positive-definite covariance matrix *Σ*, but our framework allows for more general spatial processes (*B*(*t*))_*t⩾0*_. The primary tumor stays at the same location until it dies. We assume that the shedding time *ν* is exponentially distributed, *ν* ~ Exp(*µ*) for some parameter *µ* ⩾ 0, using the convention that *V* = ∞ if *V* ~ Exp(0). Our approach to model the shedding time and other random times (introduced below) by exponential distributions is motivated by the memoryless property, which characterizes the exponential distribution: assuming that the tumor is big enough, cell clusters are shed randomly approximately at a constant rate, which implies an exponential distribution for the shedding time. Moreover, our assumption of exponentially distributed times between events (shedding and establishment) is supported biologically by the successful confrontation of Hanin et al.’s stochastic model to clinical data (Hanin et al., 2016).

After an additional random time *τ* (namely, at time *ν* + *τ*), the cell cluster settles down and forms a metastasis. We assume that *τ* is exponentially distributed, *τ* ~ Exp(*λ*) for some parameter *λ* ⩾ 0. When the cell cluster has settled down, it requires time to grow into a metastasis until it is able to shed cell clusters on its own. We model this time again with an exponential distribution with parameter *µ* ⩾ 0. Established metastasis and the primary tumor will continue to emit further cell clusters after a time which is exponentially distributed with the same parameter *µ* ⩾ 0, independently of the other random variables and processes. Cell clusters that are moving are destroyed at a rate *δ*_1_ ⩾ 0 while stationary tumors die at a rate *δ*_2_ ⩾ 0, again independently of the other particles, the movement and the growth. The process is repeated ad infinitum. Figure 1 illustrates our model.

**Fig 1.**
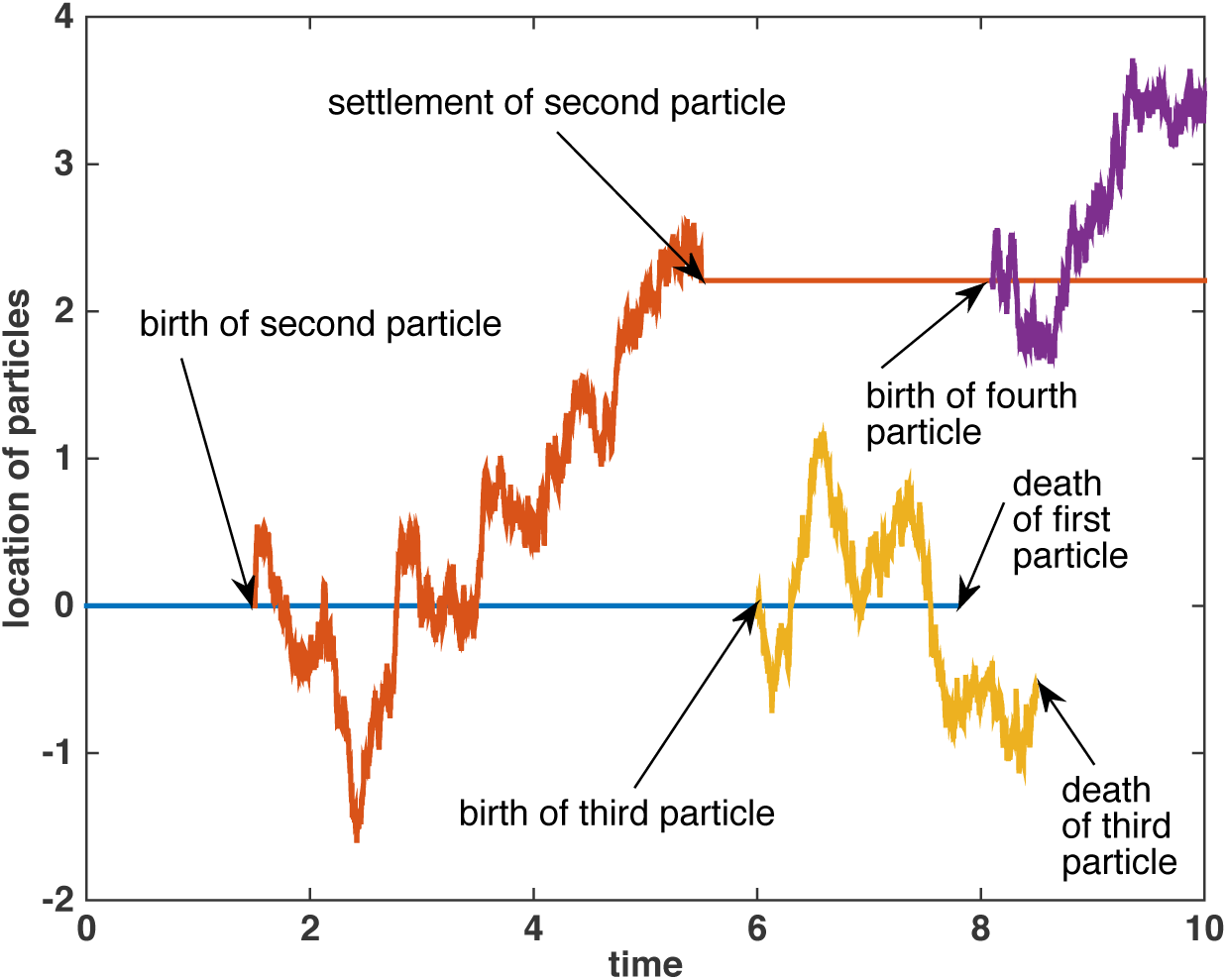
Illustration of a one-dimensional branching stochastic process with settlement. Color figure online.

A key benefit of studying a *d*-dimensional stochastic process is that we can extend the notion from cell clusters and metastases, whose movements are restricted to a system of blood and lymphatic vessels, to particles in a *d*-dimensional space. Indeed, the branching stochastic process with settlement is also relevant to other applications such as seed dispersal, epidemic spread, and forest fire spread (Martin and Hillen, 2016). In those cases, particles would refer to plant seeds, infectious agents, and burning branches, respectively. Hence “particle” can have very different meaning. In this paper, our focus is on CTCs for moving particles and metastases for stationary particles.

We denote by *N*(*t*) the number of particles born before time *t*. Their positions at time *t* are *X*_*i*_(*t*) for 1 ⩽ *i* ⩽ *N*(*t*) where we enumerate the particles by their birthdates. For fixed *t* and *i, N*(*t*) and *X*_*i*_(*t*) are random variables with values in ℕ and ℝ^*d*^, respectively. We denote by *M*(*t*) the number of particles alive at time *t*. We are interested in the following quantities:

- expected location 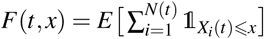: the expected number of particles located in the set (−∞, *x*^1^] ×…× (−∞, *x*^*d*^] at time *t*,
- locational variance 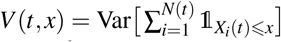: the variation in the number of particles located in (−∞, *x*^1^] ×…× (−∞, *x*^*d*^] at time *t* due to randomness,
- furthest particle distribution *H*(*t,exponentially and the asymptotic survival probabilityx*) = *P*[max_*i*=1,*…,N*(*t*)_ *X*_*i*_(*t*) *⩽ x*]: the probability that all particles are located in (−∞, *x*^1^] *×…×* (−∞, *x*^*d*^] at time *t*,
- survival probability *Q*(*t*) = *P*[*M*(*t*) > 0]: the probability that there is at least one particle alive at time *t*,

where we use the convention that 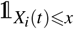 is zero for all *x* and times *t* after the death time of particle *i*. We set *H*(*t, x*) = 0 for all *x* if no more particle exists at time *t*.

### 2.1 Characterization via integro-differential equations

We use the above branching stochastic process with settlement to find a common type of equation for the expected location *F*, the variance *V*, the distribution of the furthest particle *H* and the survival probability *Q*.

#### Theorem 2.1

Consider the branching stochastic process with settlement defined above. The quantities *F,V, H, Q* satisfy

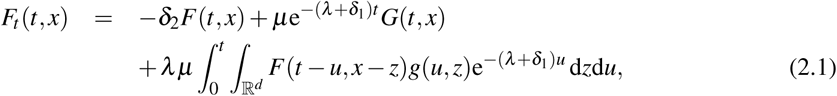

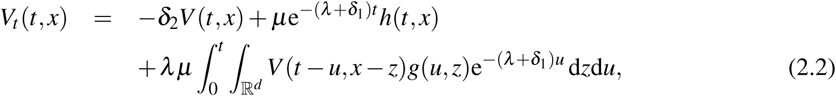

with

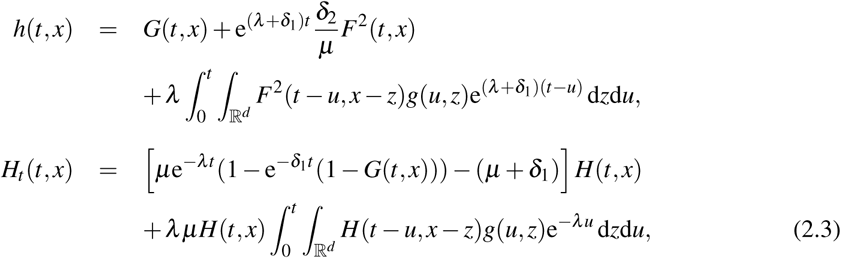

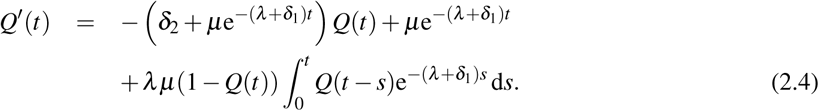

Before we present the proof of Theorem 2.1 in Section 2.3, we bring the above four equations (2.1)–(2.4) into a common form using a convolution notation. We encounter integral kernels that depend on one variable and on two variables. We use the same convolution symbol for both cases; given two kernels *k*_1_(*t, x*) and *k*_2_(*t*) and two test functions *f* (*t, x*) and *g*(*t*), we denote

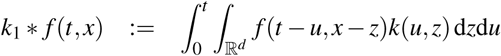

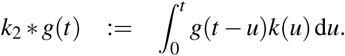

Then we combine equations (2.1)–(2.4) in the compact form

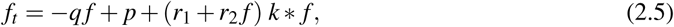

where we identify parameters and functions as shown in Table 1.

**Table 1.** Parameter values for the different cases of expected location *F*, locational variance *V*, distribution of the furthest particle *H* and survival probability *Q*, using the abbreviation 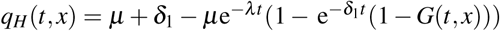.

| Case | $r_1$ | $r_2$ | $q(t, x)$ | $p(t, x)$ | $k(u, z)$ |
| --- | --- | --- | --- | --- | --- |
| $F$ | 1 | 0 | $\delta_2$ | $\mu e^{-(\lambda + \delta_1)t} G(t, x)$ | $\lambda \mu e^{-(\lambda + \delta_1)u} g(u, z)$ |
| $V$ | 1 | 0 | $\delta_2$ | $\mu e^{-(\lambda + \delta_1)t} h(t, x)$ | $\lambda \mu e^{-(\lambda + \delta_1)u} g(u, z)$ |
| $H$ | 0 | 1 | $q_H(t, x)$ | 0 | $\lambda \mu e^{-\lambda u} g(u, z)$ |
| $Q$ | 1 | −1 | $\delta_2 + \mu e^{-(\lambda + \delta_1)t}$ | $\mu e^{-(\lambda + \delta_1)t}$ | $\lambda e^{-(\lambda + \delta_1)u}$ |

In Section 3, we will analyze the general type (2.5) of non-local integro-differential equation with distributed delay and show positivity as well as local and global existence of solutions using a fixed-point argument. For the qualitative analysis, we encounter an index *R*_0_, which can be understood as the *metastatic reproduction number*:

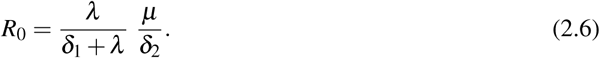

The fist factor *λ/*(*δ*_1_ + *λ*) denotes the settlement rate divided by the sum of the settlement rate and the death rate during movement. It describes the odds of being still alive while settling. The second factor in (2.6),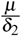, which refers to stationary particles, can be explained similarly, but here the effect of births of new particles needs to be considered. Indeed, when the ratio between shedding rate *µ* and total change rate *δ*_2_ + *µ* (stationary deaths and shedding) is summed over all potential offsprings of the particles, we obtain

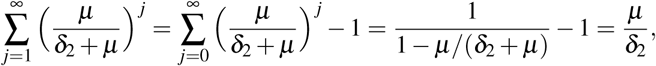

regaining the second term in (2.6).

Before we go into the details, we paraphrase the results, assuming *µ >* 0 and *λ >* 0:

- **Integro-differential equations:** *Under suitable assumptions* (*A1*), *defined later, the solutions to the above integro-differential equations* (*2.1*)*–*(*2.4*) *exist as mild solutions globally in time, whereby the two probabilities H and Q are bounded by 1. See Theorem 3.1 for global existence and uniqueness to the general type* (*2.5*) *of non-local integro-differential equations.*
- **Asymptotic properties:** *The asymptotic behavior is characterized by R*_0_:

**–** *R*_0_ < 1: *The expected number of particles shrinks exponentially and the particles die out asymptotically with certainty.*
**–** *R*_0_ > 1: *The expected number of particles grows exponentially and the asymptotic survival probability is given by* 1 − 1*/R*_0_ ∈ (0, 1).
**–** *R*_0_ = 1: *The particles die out asymptotically with certainty while their number explodes on a set with vanishing probability so that their expected number converges to a finite limit* 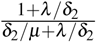. *See Lemma 3.3 for a threshold result of the survival probability Q and Corollary 3.1 for the asymptotic behavior of the expected particle number E*[*M*(*t*)].

Note that the survival probability in case *R*_0_ > 1 equals 1–1*/R*_0_, which corresponds to the critical immunization threshold in common models for infectious diseases (Hethcote, 2000). As we will explain in Section 3.2, the asymptotic properties can be seen as a compact characterization in terms of *R*_0_ for the extinction-survival dichotomy of two-type branching processes.

#### Remark 2.1

While we have assumed an exponential distribution for the shedding time, an equation for the expected location and other key quantities can also be derived considering a general distribution for the shedding time, but at a price of making the equation (and thus its potential analysis) much more complicated. Assume that the shedding times are independent and have a continuous probability distribution with density *α*. Similarly to the proof of Theorem 2.1, one can generalize (2.1) and show that the expected location *F* satisfies

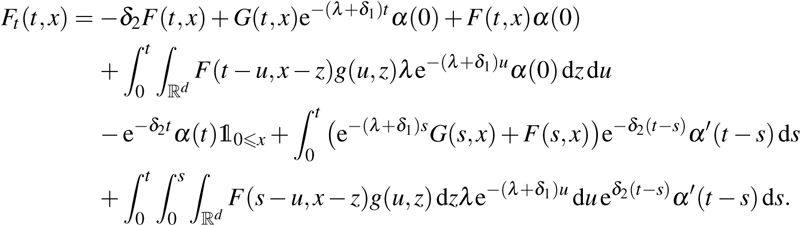

In the case of an exponential distribution, we have *α′*= −*µα* and the above two last lines equal to −*µF*(*t, x*), leading to the simpler integro-differential equation given in (2.1).

### 2.2 Instantaneous relocation

In many biological situations, the time scale of movement is very different from the time scale of reproduction or death. Most mammals, for example, explore their environment on a daily basis, while they reproduce once per year or less. And even cells in tissues can travel relative large distances in minutes, while typical cell cycles are about 8–12 h or longer. Hence on the macroscopic time scale of birth and death, relocation appears to be instantaneous. To model this situation, we consider the limit *λ* → ∞ in (2.5). When individuals are moving, they will jump immediately to a new location with density function of the distances travelled given by *g*(0,*x*). We will then ignore the time in *g*(0,*x*) and simply use a spatial relocation density *g*(*x*).

For fixed *t*, we consider a common term in (2.5)

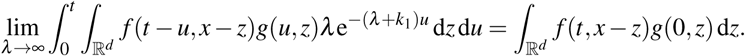

Using integration by parts yields

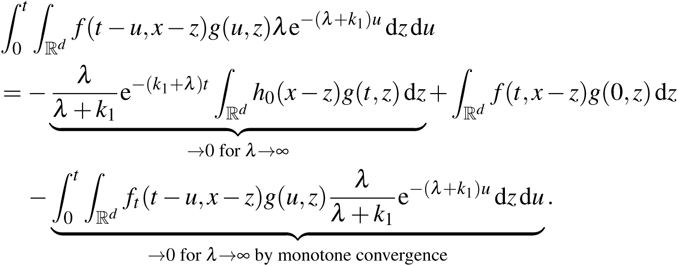

Applying this scaling to the four equations (2.1)–(2.4) from Theorem 2.1 and using *g*(*x*) instead of *g*(0, *x*), we find

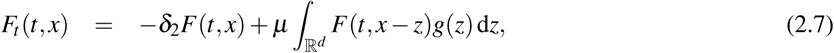

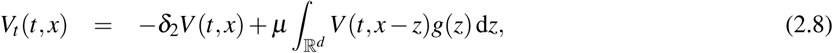

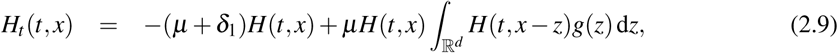

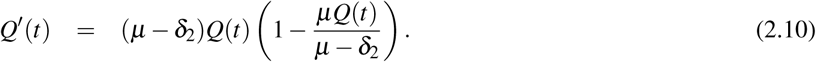

The equations for the instantaneous relocation case (2.7)–(2.9) are linear integral equations as they have been studied in the classical literature (Krasnoselskii, 1964). They are also of the form of *birth-jump processes* as introduced by Hillen et al. (2015). Hence the branching stochastic process with settlement appears as a generalization of birth-jump processes.

The metastatic reproduction number (2.6) becomes in this case

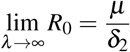

The survival probability *Q* satisfies a logistic differential equation (2.10), which can be analyzed quickly. If *R*_0_ < 1, then the origin is asymptotically stable, and solutions converge to 0: metastasis dies out. If *R*_0_ > 1, then *Q* converges to a limit

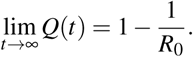

If *R*_0_ = 1, then *Q*(*t*) = *Q*(0) is constant.

This also corresponds to the classical result for the asymptotic extinction probability of a birth-death process; see, for example, Section 4.4 of Durrett (2012). Indeed, in the limit *λ* → ∞, the number of particles evolves according to a Markov chain starting at 1 and with jump rates *q*(*i,i* + 1) = *µi* and *q*(*i,i*− 1) = *δ*_2_*i* when there are *i* particles.

### 2.3 Derivation of the integro-differential equations (proof of Theorem 2.1)

The main idea to derive equations (2.1)–(2.4) from Theorem 2.1 is to use a renewal argument based on the recursive structure of branching stochastic processes with settlement. Since this goes similarly for the different functions, *F,V,H*, and *Q*, we give a detailed proof in the case of *F* and mention afterwards how to start the proofs for *V*,*H* and *Q*, with the subsequent analysis being done analogously to that of *F*. We start by conditioning on time *V* of the birth of the second particle, which means writing

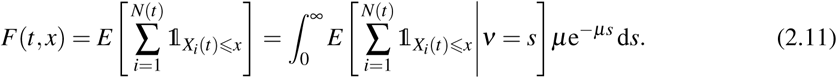

In the event {*V* > *t*}, the birth of the second particle happens only after time *t* so that at time *t* either only the first particle exists, which is located at zero, or no particle at all exists if the first particle has died before time *t*. We denote by *A*_2_(*t*) the event that the first particle has not died before time *t*, which has probability 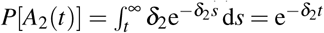. For *s* > *t*, we have

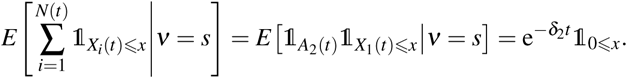

On the other hand, in the event {*V* ⩽ *t*}, the second particle was born before time *t* if the first particle was then alive. We need to distinguish two cases on {*V* ⩽ *t*}. The first case is when the second particle has not yet settled down at time *t*. This happens when *V* + *τ* > *t* and then the distribution of 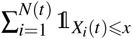 conditional on *V* = *s* and the first particle being alive at time *s* is equal to the sum of 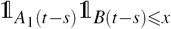 and 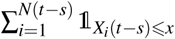, where *B*(*t-s*) is an independent stochastic process at time *t-s* (movement of second particle) with *A*_1_(*t-s*) the event that the second particle born at time *s* is alive at time *t*. The term 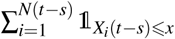 arises because we can start again at time *V* = *s* in the same way as at time zero thanks to the recursive structure and the memoryless property. The probability of the first particle being alive at time *s* is 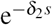 while the survival probability of the second particle is

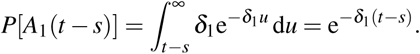

Therefore, we have for *s* ⩽ *t* and *u* > *t−s* that

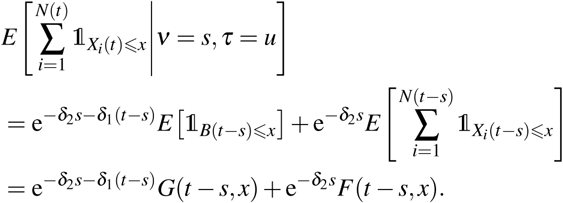

For the other case where *V* + *τ* ⩽ *t*, the second particle has already settled down before time *t* and behaves in the same way at *V* + *τ* as the first particle did at time 0, but with starting point *B*(*τ*) and conditional on being then still alive. Therefore, the distribution of 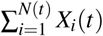 conditional on *V* = *s, τ* = *u* and the first particle being alive at time *s* is then equal to the sum of independent copies of the random variables 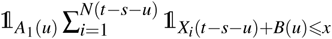 and 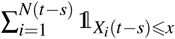. This yields for *s* + *u* ⩽ *t* that

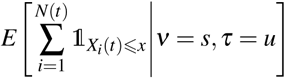

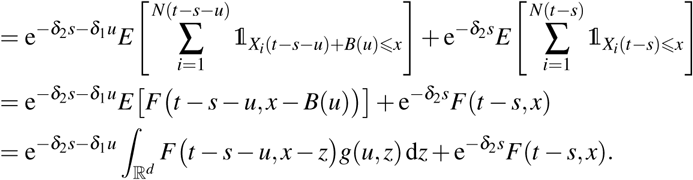

Combining the terms, we obtain

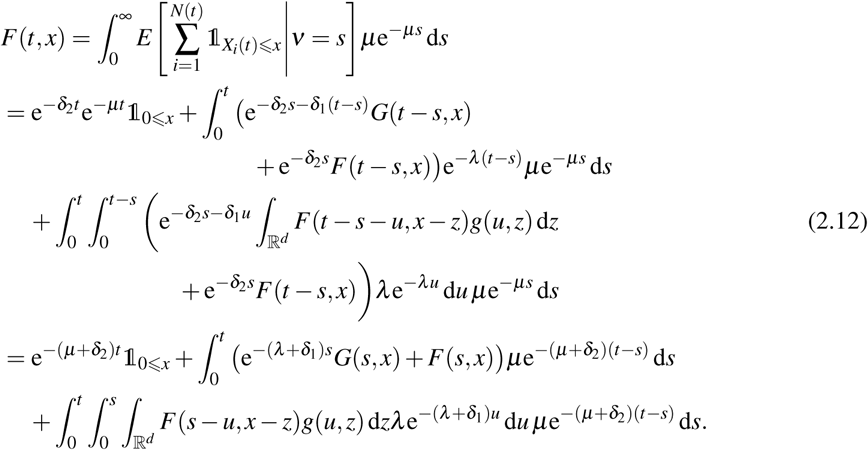

We deduce by differentiating with respect to *t* that

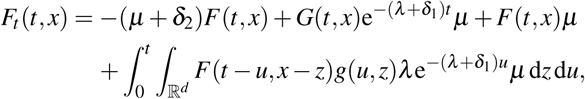

which shows that *F* satisfies (2.5) for the choices mentioned in Table 1.

The proof for the integro-differential equations of *V*, *H* and *Q* go analogously. For *V*, we start by decomposing it as

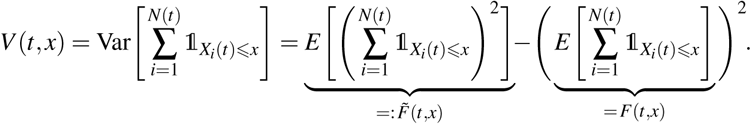

This implies 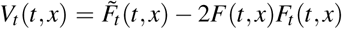 so that the integro-differential equation of *V* follows from such equations for *F* and 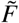. Similarly to (2.11), we write

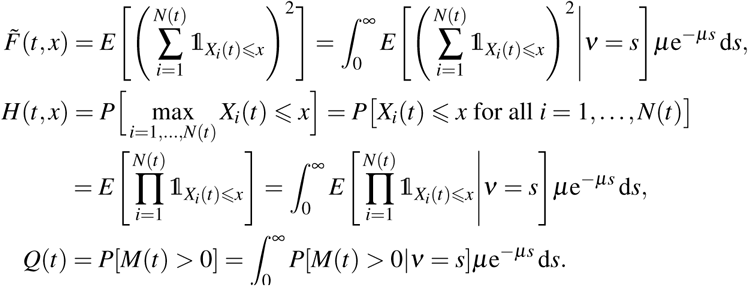

The subsequent steps for 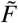, *H* and *Q* go analogously to the analysis done for *F* by using decompositions similar to (2.12).

## 3. Analysis of the integro-differential equations

In this section, we first prove existence and uniqueness results for the integro-differential equations (2.5). We then analyze the mean numbers of moving and stationary particles, where we already see that *R*_0_ has a threshold of 1 that distinguishes between metastatic growth or decay. This result for *R*_0_ is derived in two ways: from a stability analysis of differential equations and, alternatively, from general results for two-type branching processes.

### 3.1 Local and global existence

For the general type (2.5) of integro-differential equations, we make the following assumptions:

**Assumptions (A1):**

- We assume *r*_1_ ⩾ 0, *r*_2_ ∈ ℝ,
- *q*(*t, x*) ⩾ *δ* > 0 is uniformly bounded and Lipschitz continuous in *t* and *x*.
- For each *T* > 0 *p*(*t, x*) ⩾ 0 is uniformly bounded on [0, *T*] by *P*, absolute continuous in *x* and continuous in *t*.
- For the cases *F, H,V*: *k*(*u, z*) ⩾ 0 is continuous in *x* and satisfies for some constant *K* > 0 that

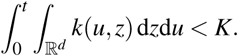
- For the case *Q*: *k*(*u*) ⩾ 0 satisfies for some constant *K* > 0 that

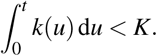
- The initial condition *f*_0_(*x*) = *f* (0, *x*) ⩾ 0 is bounded in *L*^∞^ (ℝ^*d*^).

#### 3.1.1 Mild solutions

To prove local existence and uniqueness, we consider *T* > 0 and use as phase space

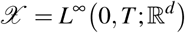

with the essential supremum norm factor || · ||_∞_. To find a mild formulation of (2.5), we define an integrating factor

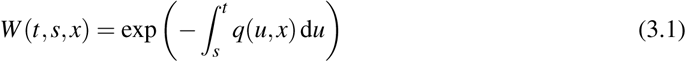

where the *x*-dependence arises only in the case of *f* = *H*. Since *q* is Lipschitz continuous and *q* > 0 we have that *W* (*t, s, x*) is a non-negative evolution family with 0 < *W* (*t, s, x*) ⩽ 1 for all 0 ⩽ *s* ⩽ *t* ⩽ *T*. We use the variation of constant formula to formally solve (2.5) as

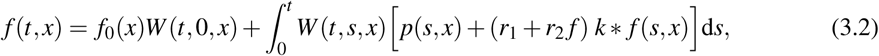

with *f* (0, *x*) = *f*_0_(*x*), which is our mild formulation.

Based on the various interpretations of *f* as probability or variance, we are interested in non-negative solutions.

##### Lemma 3.1

Assume (A1) and let *f* ∈ 𝒳 be a mild solution of (2.5).

1. Then *f* (*t, x*) ⩾0 as long as the solution exists.
2. If *f* (*t*_0_, *x*_0_) = 0 for a point (*t*_0_, *x*_0_), then this implies that

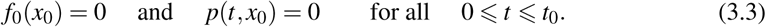
3. If in addition *p*(*t, x*) > 0 for all (*t, x*) ∈ (0, *T*) *×* ℝ^*d*^ then *f* (*t, x*) > 0 for all (*t, x*) ∈ (0, *T*) *×* ℝ^*d*^.

*Proof.*

1. Consider any time *t*_0_ such that *f* (*t, x*) ⩾0 for all 0 ⩽ *t* ⩽ *t*_0_ and all *x* ∈ ℝ^*d*^. Define

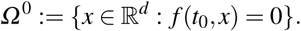 Then at each of the points (*t*_0_, *x*), *x* ∈ *Ω* ^0^, we have the following signs of the terms in (2.5):

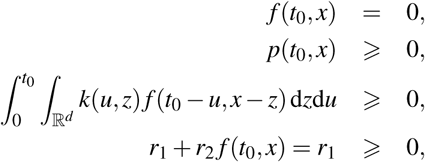

hence from (2.5), we find 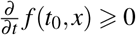 for all *x* ∈ *Ω* ^0^, and the solution does not decay further.
2. Taking a point (*t*_0_, *x*_0_) with *f* (*t*_0_, *x*_0_) = 0, we find from the mild formulation (3.2) that

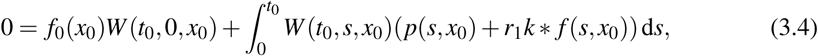

where all terms are non-negative. Hence (3.3) follows.
3. If *p* > 0, we see that (3.4) is not possible, hence *f* > 0.

##### Proposition 3.1

Assume (A1). Then there exists a time *T* > 0 and a unique mild solution *f* ∈ 𝒳 which satisfies (3.2).

*Proof.* Given a function 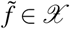, we define an iteration operator 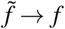 where *f* is the unique solution of

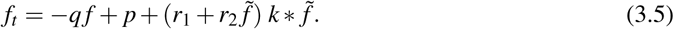

Using the integrating factor (3.1) we solve (3.5) as

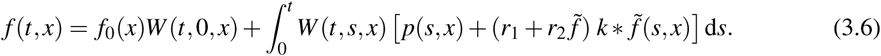

**Step 1:** Given an initial condition *f*_0_ ∈ 𝒳 with norm ‖ *f*_0_‖ _𝒳_ = *M*_0_. We show that for each constant *M* > 0 there exists a time *T*_1_ > 0 such that *A*: *B*_*M*_(*f*_0_) *→ B*_*M*_(*f*_0_) for all *T* ⩽ *T*_1_, where *B*_*M*_(*f*_0_) ⊂ 𝒳 denotes the closed ball of radius *M* in 𝒳 with center *f*_0_. Consider 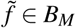 then 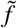 has norm

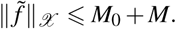

From (3.6) we find

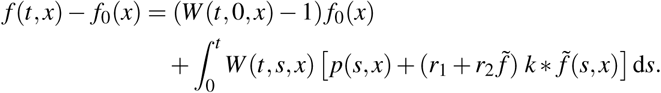

Since *W* (*t*, 0, *x*) < 1 for all *t* > 0, we can ignore the first term and estimate

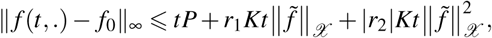

which implies that

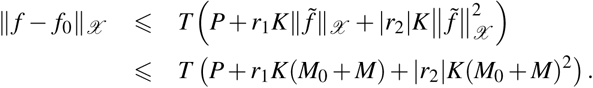

To ensure ‖ *f – f*_0_‖_𝒳_ ⩽ *M*, we need

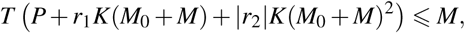

which is true for all

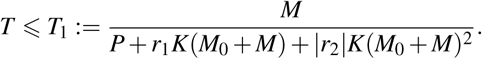

Hence, for *T* ⩽ *T*_1_, we have *A*: *B*_*M*_(*f*_0_) *→ B*_*M*_(*f*_0_).

**Step 2:** We show that *A* is a contraction for *T* ⩽*T*_2_ for some *T*_2_ > 0.

Given 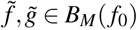 with 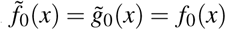. Then from (3.6), we find

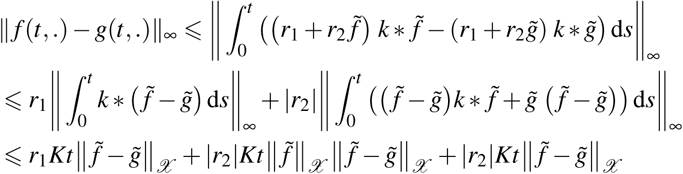

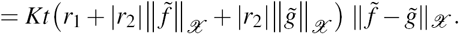

Hence

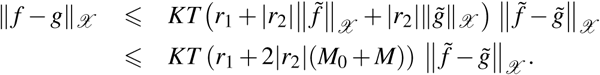

Then *A* is a contraction for all

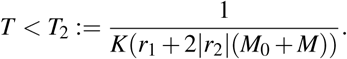

**Step 3:** For *T* < min {*T*_1_, *T*_2_} the Banach fixed point theorem applies, and *A* has a unique fixed point in 𝒳, which satisfies the mild formulation (3.2).

The local existence times *T* from the above proof depend on the norm of the initial condition, hence we cannot simply repeat the argument to obtain global existence. Nevertheless, we show next that solutions are global.

#### 3.1.2 Global existence

##### Theorem 3.1

The unique mild solutions from Proposition 3.1 exist for all times. The probabilities *H* and *Q* are globally bounded by 1 (as solutions of their corresponding integro-differential equation).

*Proof.*

- For the cases *F* and *V* we have *r*_2_ = 0, hence (2.5) becomes a linear equation in *f*. We estimate

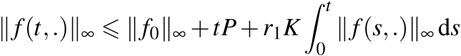

and Gronwall’s lemma implies that ‖*f* (*t*,.)‖_∞_ grows at most exponentially, hence proving global existence.
- In the case *Q* the parameter *r*_2_ =−1 is negative. Hence in the global estimate, we can remove the quadratic term. Again Gronwall’s lemma applies and global existence follows. To show that the survival probability *Q* is bounded by 1, we assume that *Q*(*t*_0_) = 1 and *Q*(*t*) < 1 for all 0 ⩽ *t* < *t*_0_, or *t*_0_ = 0 and *Q*(0) = 1. Then at *t*_0_ we find from (2.4) that

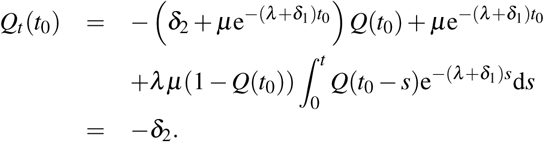 Hence *Q* decays near 1. In our specific application we have *Q*(0) = 1, hence *Q* decays initially and as survival probability, *Q*(*t*) is non-increasing.
- A similar argument is applied for *H*. We assume that *H*(*t, x*) ⩽1 for all *t* ⩽ *t*_0_ and that there exists a point *x*_0_ with *H*(*t*_0_, *x*_0_) = 1. Then at (*t*_0_, *x*_0_) we observe

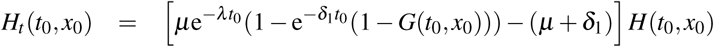

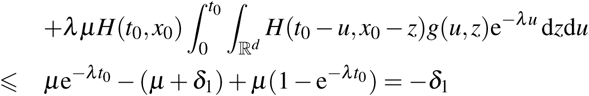

and *H*(*t*_0_;*x*_0_) = 1 decays.

### 3.2 Metastatic reproduction number R_0_

To introduce the metastatic reproduction number, we reduce the above model by looking at the expected numbers of moving and stationary particles. We denote by *a*(*t*) and *b*(*t*) the expected numbers of moving and stationary particles, respectively. Moving particles die at rate *δ*_1_ and become stationary at rate *λ* while new moving particles are born at rate *µ* from the stationary particles. This reasoning leads to

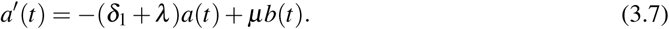

Formally, we can prove (3.7) analogously to the first part of the proof of Theorem 3.2. Similarly, stationary particles die at rate *δ*_2_ and moving particles become stationary at rate *λ*, leading to

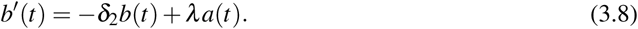

The differential equations for *a* and *b* form a linear system with coordinate matrix

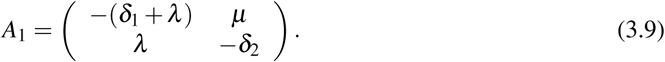

The matrix *A*_1_ has trace and determinant as

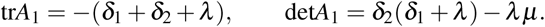

The trace is negative, hence the origin is asymptotically stable for det*A*_1_ > 0 and unstable for det*A*_1_ < 0. Using

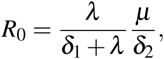

we find the following result:

#### Lemma 3.2

Consider (3.7), (3.8).

- If *R*_0_ < 1 then (0, 0) is globally asymptotically stable.
- If *R*_0_ > 1 then (0, 0) is unstable (it is a saddle).
- If *R*_0_ = 1 then (0, 0) is non-hyperbolic and we have a continuum of steady states in direction (*δ*_2_, *λ*)^*T*^.

*Proof.* For the first two statements note that we can write

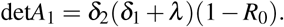

For the third statement we find that for *R*_0_ = 1 the vector (*δ*_2_, *λ*)^*T*^ is an eigenvector of *A*_1_ with eigenvalue 0.

It should be noted that *a*(*t*)+ *b*(*t*) = *E*[*M*(*t*)] with the formula for the expected total number *E*[*M*(*t*)] of particles.

Using specific initial conditions (*a*(0), *b*(0)) = (0, 1) we can explicitly solve equations (3.7), (3.8) and we find the asymptotic ratio of moving versus stationary particles

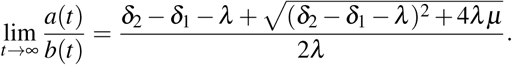

An alternative way to analyze the asymptotic number of particles is to relate our process to a two-type branching process in discrete time and to use results available for such processes. The two types are the moving and stationary particles. As explained in Section 2.3 of (Haccou et al., 2005) and Section 6.2 of (Kimmel and Axelrod, 2015), for multi-type branching processes, the asymptotic extinction-survival behavior is characterized by the Perron root *ρ* (maximal eigenvalue that corresponds to an eigenvector with positive entries) of the mean matrix *M*, which generalizes the mean reproduction in the one-type case. The process is called subcritical if *ρ <* 1, critical if *ρ* = 1, and supercritical if *ρ >* 1. For our model, the mean matrix is given by

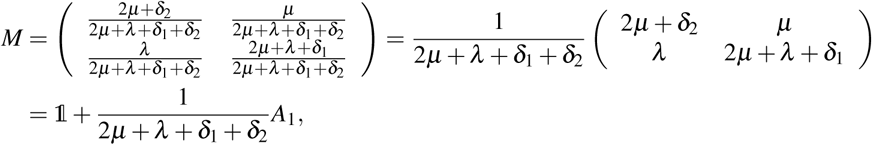

where we recall *A*_1_ from (3.9). If *v* is eigenvector of *A*_1_ with eigenvalue *α*, then it is eigenvector of *M* with eigenvalue 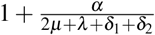. Hence, *M* has an eigenvalue greater than 1 if and only if *A*_1_ has a positive eigenvalue, which is the case if and only if *R*_0_ > 1, as explained before Lemma 3.2. Also note that such an eigenvalue corresponds to an eigenvector

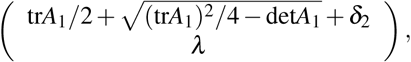

which has positive entries. Therefore, the Perron root *ρ* of the mean matrix *M* is greater than one (supercritical case) if and only if *R*_0_ > 1. Similarly, the critical case *ρ* = 1 and the subcritical case *ρ <* 1 are equivalent to *R*_0_ = 1 and *R*_0_ < 1, respectively.

### 3.3 Survival probability Q(t)

The survival probability *Q* is a special case of (2.5), where there is no spatial variable. In this case, (2.5) becomes

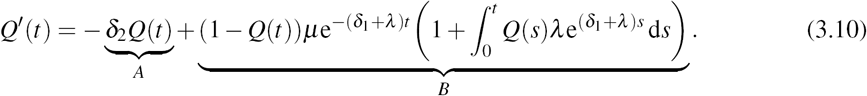

where

A: death rate of original particle times the survival probability at time *t*. Note that this is the only term if *µ* = 0 (no births), in which case, the survival probability equals *Q*(*t*) = exp(−*δ*_2_*t*).

B: correction term because the birth of particles leads to a higher survival probability than in the case *µ* = 0. If the original particle has offsprings, all of the original particle, the offsprings of the original particle and further offsprings must die to extinct all particles, which is reflected in the term B.

Based on (3.10), we can derive a second-order ODE for *Q*, namely,

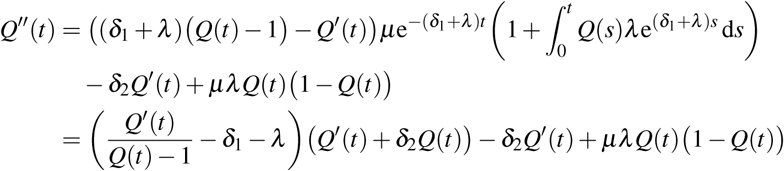

This equation can be transformed into a system of first-order ODEs as

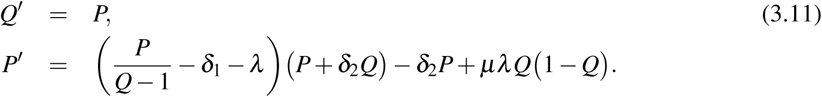

We show the following result:

#### Lemma 3.3

Consider system (3.11).

- If *R*_0_ < 1 then (0, 0) is locally asymptotically stable.
- If *R*_0_ > 1 then (0, 0) is locally unstable (it is a saddle).

*Proof.* The linearization of (3.11) at (0, 0) is

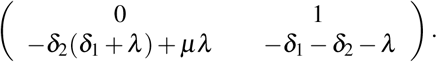

This matrix has the same trace and determinant as the matrix *A*_1_ that we encountered in Lemma 3.2. Hence for *R*_0_ < 1, the origin is locally asymptotically stable, and for *R*_0_ > 1, it is a saddle.

### 3.4 Asymptotic behavior as t → ∞

In this section, we study how the average number of particles and the survival probability behave in the limit *t →* ∞. We first provide an explicit formula for the average number of particles at an arbitrary time *t*.

#### Theorem 3.2

For *µ* ≠ 0, and *λ* ≠ 0 or *δ*_1_ ≠ *δ*_2_, the average number of particles alive at time *t* is given by

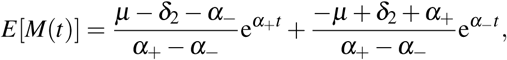

where

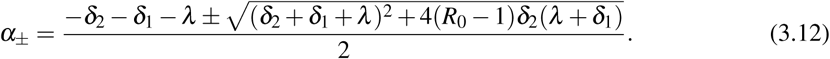

For *µ* = 0 or both *λ* = 0 and *δ*_1_ = *δ*_2_, the average number of particles at time *t* equals 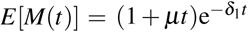.

*Proof.* Similarly to the proof of Theorem 2.1, we find

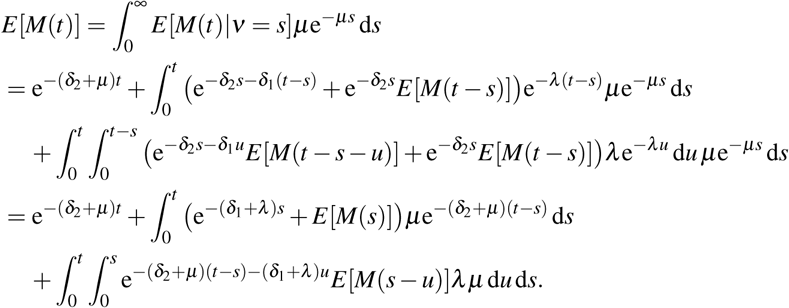

We deduce that

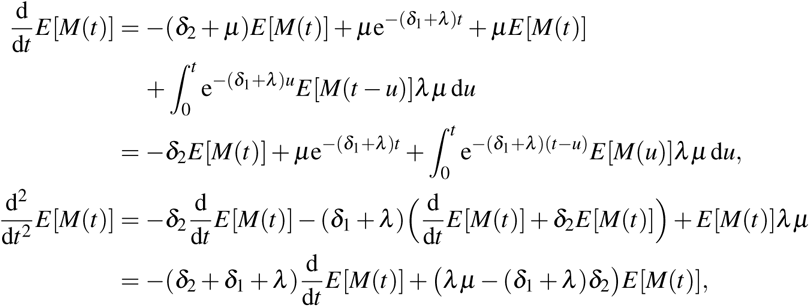

which shows that *E*[*M*(*t*)] satisfies a linear ordinary differential equation, whose solution is of the form 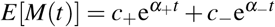 with

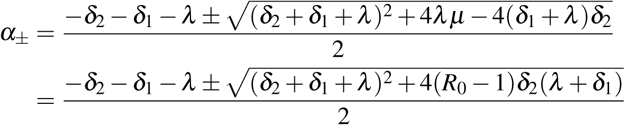

if *δ*_2_ ≠ *δ*_1_ + *λ* or *λ µ* ≠ 0. To find the constants *c*_−_ and *c*_+_, we use the boundary conditions

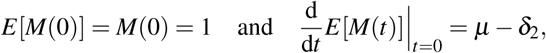

where we applied that at time 0 there is one particle which will be either eliminated (at rate *δ*_2_) or doubled (at rate *µ*). This yields

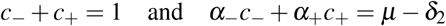

which gives 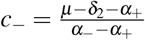 and 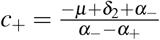. In the case *λ* = 0 and *δ*_1_ = *δ*_2_, we have

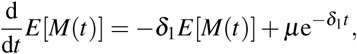

which satisfies

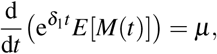

hence 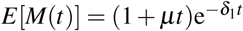. In the case *µ* = 0, we have 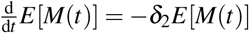 so that 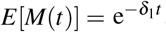.

For *µ* ≠ 0, and *λ* ≠ 0 or *δ*_1_ ≠ *δ*_2_, we obtain from Theorem 3.2 that

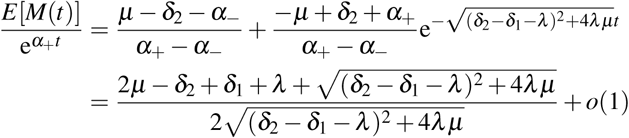

for *t →* ∞ so that *E*[*M*(*t*)] has exponential growth with rate *α*_+_.

Note that *α*_+_ > 0 in (3.12) if and only if *R*_0_ > 1 and *α*_+_ < 0 whenever *R*_0_ < 1. Also note that *α*_+_ is real for all *R*_0_ ∈ [0, 1]. From this reasoning, we deduce the following result.

#### Corollary 3.1

*E*[*M*(*t*)] has the following asymptotic behavior: it …

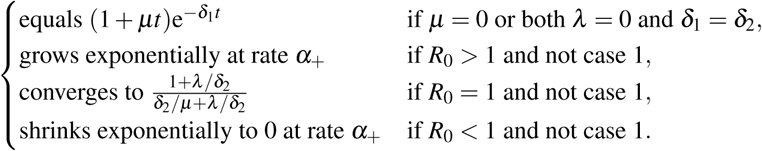

In Lemma 3.3 we have already shown the corresponding threshold phenomenon for the survival probability *Q*. In more detail, we can also compute the asymptotic value as *t →* ∞.

#### Lemma 3.4

The asymptotic survival probability is

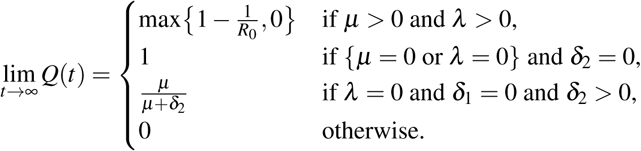

*Proof.* We first observe that if *δ*_2_ = 0, then *Q*(*t*) = 1 for all *t* and hence lim_*t→*∞_ *Q*(*t*) = 1 because the first particle then lives forever. Now consider the case where *δ*_2_ > 0 and *δ*_1_ = *λ* = 0 so that (3.10) becomes

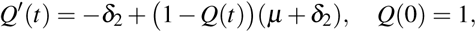

whose solution is 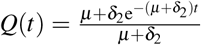, which converges to 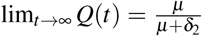. Let us now consider *δ*_1_ + *λ >* 0 and *δ*_2_ > 0. Because *M*(*t*) = 0 implies *M*(*s*) = 0 for all *s ⩾ t* (after particles died out, there will always be zero particles), we obtain

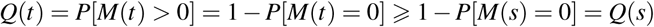

for all *s ⩾ t*, which shows that *Q* is a non-increasing function. Since *Q* is also bounded from below by 0, it converges to some limit *Q*(∞):= lim_*t→*∞_ *Q*(*t*). Using lim_*t→*∞_ *Q*^*’*^(*t*) = 0 and taking limits in (3.10) gives

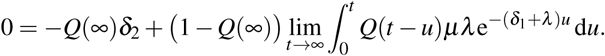

From L’Hôpital’s rule, it follows that

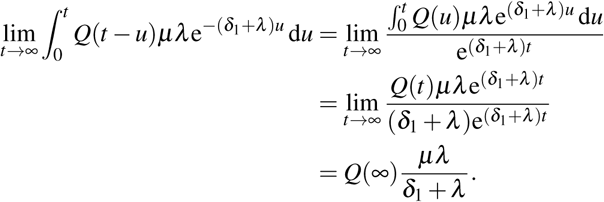

Therefore, *Q*(∞) satisfies the quadratic equation

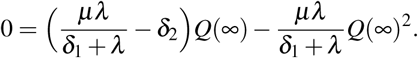

If *λ* = 0 or *µ* = 0, its solution is *Q*(∞) = 0 while otherwise, the solutions are zero or

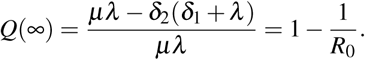

Note that *Q*(*t*) cannot cross the level 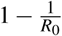 so that it will converge at 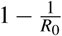 if this quantity is greater than 0.

## 4. Murine Metastatic Cancer Data

In this section, we apply the modeling framework developed above in the context of metastatic cancer data for mice. We recall that moving particles correspond to CTCs while stationary particles correspond to metastases. This interpretation requires an appropriate identification of the model parameters from biologically reasonable estimates obtained from the literature. We then make a brief note on the numerical implementation, followed by a presentation of the results themselves.

### 4.1 Murine data

Because of the scarcity of quantitative data for metastatic cancer in humans, the majority of the values discussed below have come from experimental models of metastasis in mice. Many of the studies mentioned below follow a similar procedure (those in Cameron et al. (2000); Fidler (1970); Luzzi et al. (1998); Sindelar et al. (1975) for example), and we provide a brief outline of their methods. Tumor-free mice are injected with radio-labeled cancer cells (B16 melanoma (Fidler, 1970), M19 Fibrosarcoma (Sindelar et al., 1975), B16F1(0) melanoma (Cameron et al., 2000; Luzzi et al., 1998)), and observed and/or sacrificed at various time points ranging from 1 minute to 14 day post-injection. Organs of interest (multiple organs (Fidler, 1970), lungs (Cameron et al., 2000; Sindelar et al., 1975), and liver (Luzzi et al., 1998) are removed and analyzed for the number and location of cancer cells, cancer cell clusters, and metastases. In addition to the radio-labeled cancer cells, the Chambers group (Cameron et al., 2000; Luzzi et al., 1998) injected inert microspheres that become lodged within the microvasculature of the target organ in order to accurately determine the change in cell numbers over time. Details of specific experimental models can be found in the cited references.

### 4.2 Modeling metastatic cancer

In order to apply our model to the metastatic dissemination of cancer, we must first carefully define what is meant by ‘stationary’ and ‘mobile’ particles in this context. ‘Stationary’ particles will play the role of established tumors capable of shedding mobile particles without exhausting themselves. ‘Mobile particles’, therefore, will represent small clusters of individual cancer cells that are actively circulating through the vasculature. This interpretation necessitates different scales for the two classes of particles, with established tumors consisting of at least 10^8^ cells — corresponding to a tumor volume of approximately 1cm^3^ (Del Monte, 2009) — and CTCs consisting of anything between a single cell to several dozen (Friedl and Mayor, 2017). Such a distinction requires careful attention when parameterizing the model. Below we discuss our approach to address this concern.

First we consider the shedding rate, *µ*. Assuming that an established tumor consists of 10^8^ cells (Del Monte, 2009) and that the number of CTCs shed per day range between 0.0001% – 0.01% of the cells available within the established tumor (Weiss, 1990), we may choose *µ* ∈ [100, 10000] particles per day. For the simulations presented herein, we chose *µ* = 346 particles/day. This choice was made in order to have an average of 10 established tumors by the end of 14 days (Sindelar et al., 1975) (determined using the asymptotic expected ratio of moving to stationary particles from the comment after Lemma 3.2).

Second, we estimate the stationary particle death rate, *δ*_2_. With the interpretation of a stationary particle as an established tumor, *δ*_2_ corresponds to the rate of spontaneous tumor remission. We use Jessy’s estimate of *p* = 10^−5^ (Jessy, 2011) for the probability of spontaneous remission and assume that *p* = *δ*_2_*/λ* to obtain the value of *δ*_2_ reported in Table 2.

**Table 2.** Model Parameters and the values used in presented simulations. See text for further details.

| Parameter | Description | Value | Units | References |
| --- | --- | --- | --- | --- |
| $\mu$ | Shedding rate | 346 | particles/day | Del Monte (2009);<br>Weiss (1990) |
| $\lambda$ | Establishment rate | 0.0035 | particles/day | Luzzi et al. (1998) |
| $\delta_1$ | Mobile death rate | 7 | particles/day | Chambers et al. (2002);<br>Fidler (1970);<br>Liotta and DeLisi (1977) |
| $\delta_2$ | Stationary death rate | $3.46 \times 10^{-8}$ | particles/day | Jessy (2011) |

Third, we consider the rate of mobile particle settlement, *λ*. Experimental murine models of metastasis suggest that nearly 80% of the CTCs shed from the primary tumor into the vasculature will survive through the circulation and successfully extravasate at a secondary site (Cameron et al., 2000; Luzzi et al., 1998). For the purposes of our model however, successful extravasation alone does not represent ‘settlement’. ‘Settlement’ in our model includes not only successful extravasation at a secondary site, but survival and growth to a palpable secondary tumor as well. For this reason, we use the more suggestive terminology ‘establishment’ instead of ‘settlement’. Moreover, we assume that the establishment rate, *λ*, is related to the shedding rate, *µ*, via *λ* = *µq* where *q* denotes the probability (per cell) of establishment. The probability *q* has been estimated by several investigators to range between 0.0001 and 0.00001 (Chambers et al., 2002; Fidler, 1970; Luzzi et al., 1998; Sindelar et al., 1975). In the results presented below, we have used the lower estimate of *q* = 0.00001 (Luzzi et al., 1998).

Fourth, we require an estimate for the mobile particle death rate, *δ*_1_. While approximately 80% of cancer cells released into circulation will survive in the vasculature and successfully extravasate at a secondary location (Cameron et al., 2000; Luzzi et al., 1998), the fraction of these extravasated cells that will grow and form a metastasis is very small (Cameron et al., 2000). Consequently, our estimate for the mobile particle death rate must also include the death rate of successfully extravasated cells that do not become metastases, and will be much larger than if we included only deaths during transit. Additionally, the time that CTCs spend traveling through the circulation has been estimated to be between 1 and 3 hours (Luzzi et al., 1998; Sindelar et al., 1975). Therefore, assuming that ‘mobile particle’ means ‘CTC’, we expect these particles to be short-lived. Combining these observations, and based on previous results (Chambers et al., 2002; Fidler, 1970; Liotta and DeLisi, 1977), we chose to have 99.9% of all the CTCs that are shed over the course of a day perish that day. Under this assumption, the mobile death rate becomes *δ*_1_ = 7 particles/day.

Finally, we must choose a stochastic process, (*B*(*t*))_*t⩾*0_, to model the movement of mobile particles. In the simulations presented below, we have modeled the movement of mobile particles as a scaled one-dimensional Brownian motion of the form

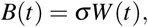

where (*W*_*t*_)_*t⩾*0_ is a standard Brownian motion, and 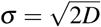 is the volatility, with *D* > 0 the *effective* diffusivity — that is to say, the diffusivity estimated assuming that movement between tumor sites is accomplished entirely by diffusion. In order to simulate the worst case scenario of full metastatic colonization of a 16cm mouse after 14 days we choose *D* = 37cm^2^/day (see Figure 4). Whereas we recognize that the transport of CTCs from a primary site to a distant secondary site through the vascular system does not occur exclusively through diffusion, we have, for the sake of simplicity, chosen to model vascular transport as a diffusive process. We leave as future work the development and inclusion of more realistic models of spatial spread based on blood circulation mechanisms (see Section 5 for further discussion).

### 4.3 Numerical results

Simulation of the model was done using a variant of the Gillespie algorithm (Erban et al., 2008; Gillespie, 1976). The key feature of this algorithm is its non-uniform time stepping which is ideally suited for our model, as our model can see (for example) the total rate of mobile particle creation double upon successful establishment of a secondary tumor because of the assumption of independence amongst the stationary particles. Our implementation of Brownian motion is equivalent to a discrete-time random walk with step sizes normally distributed with mean zero and variance 2*DΔt*, where *Δt* > 0 is the time step. For details concerning the simulation of Brownian motion using the Gillespie approach, consult (Erban et al., 2008). The implemented algorithm assumes a finite spatial domain, whereas the theoretical work presented in the previous sections does not. In order to simulate an infinite domain, we have chosen the finite domain to be ‘sufficiently large’ so that there are no collisions between our mobile particles and the domain boundaries within the time of interest. The meaning of ‘sufficiently large’ depends on the movement of the mobile particles. With the modeling choices outlined above, the spatial domain [−75cm, 75cm] divided into *K* = 1500 bins each of width 0.1cm was ‘sufficiently large’ for our purposes, as demonstrated by the fact that no individual realization had particle-boundary collisions.

In Figures 2–4, we present the average results of 1000 distinct realizations of the stochastic model simulated over a period of 14 days. Validation of our simulations is done in Figure 2 by comparing the simulated results to the exact analytic solution derived in Theorem 3.2. The average percent error over the 14 days simulated is 0.77%, with a maximum value of 2.43%. Further validation of the simulation framework was done using the extinction probability, 1 − *Q*(*t*), where *Q*(*t*) is the survival probability described by the ODE system (3.10), in which good agreement between the theoretical and empirical trajectories was obtained (results not shown). We note that we have not compared the spatio-temporal simulation results (Figures 3 and 4) to the integro-differential equation descriptions (2.1)–(2.3) due to the difficulty in accurately simulating such models, and the development of efficient numerical methods to accurately solve this type of equation is left as future work.

**Fig 2.**
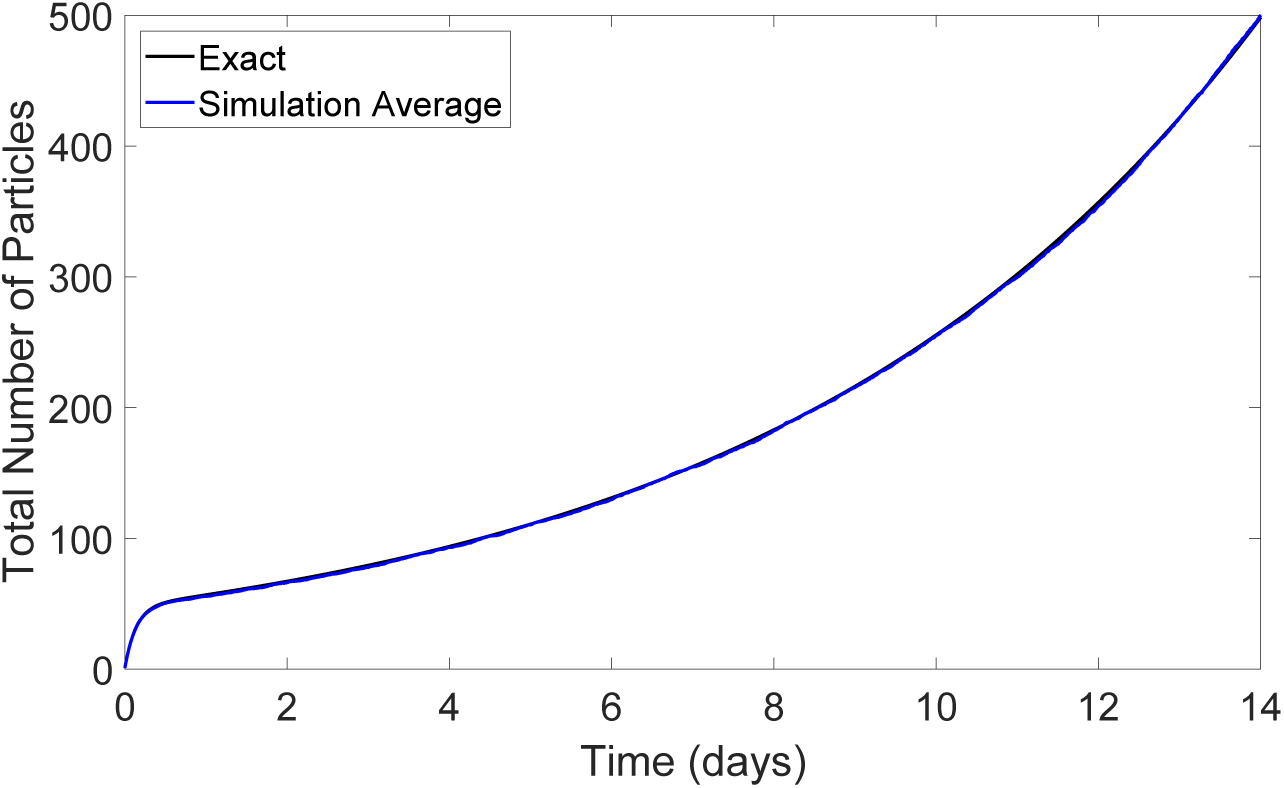
Total number of particles (stationary and mobile) as a function of time. Comparison of the theoretical expected value from Theorem 3.2 (black) and the average of 1000 realizations of the stochastic model (blue). Parameters as in Table 2. Color figure online.

**Fig 3.**
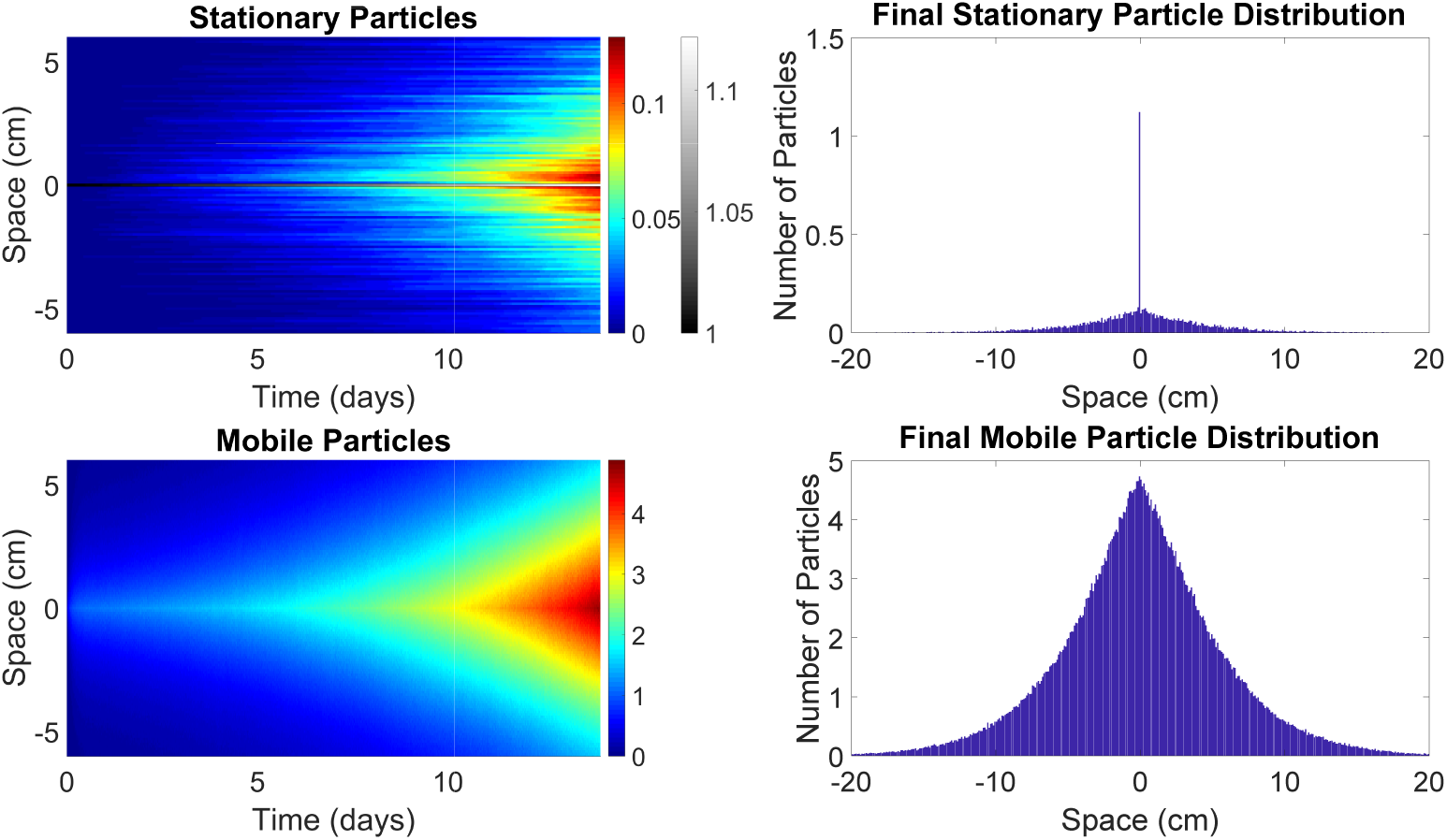
Average results of 1000 individual realizations of the stochastic model. Left column: spatio-temporal dynamics of the stationary (top) and mobile (bottom) particles. The horizontal axis is time (in days) while the vertical axis denotes space (in m from location of primary tumor). Number of particles indicated by the coloring (color figure online). Note the different scales in the top plot. Right column: average spatial distribution of the stationary (top) and mobile (bottom) particles at the end of the 14 day simulations. Note the difference in scales from top to bottom. Parameters used as in Table 2.

**Fig 4.**
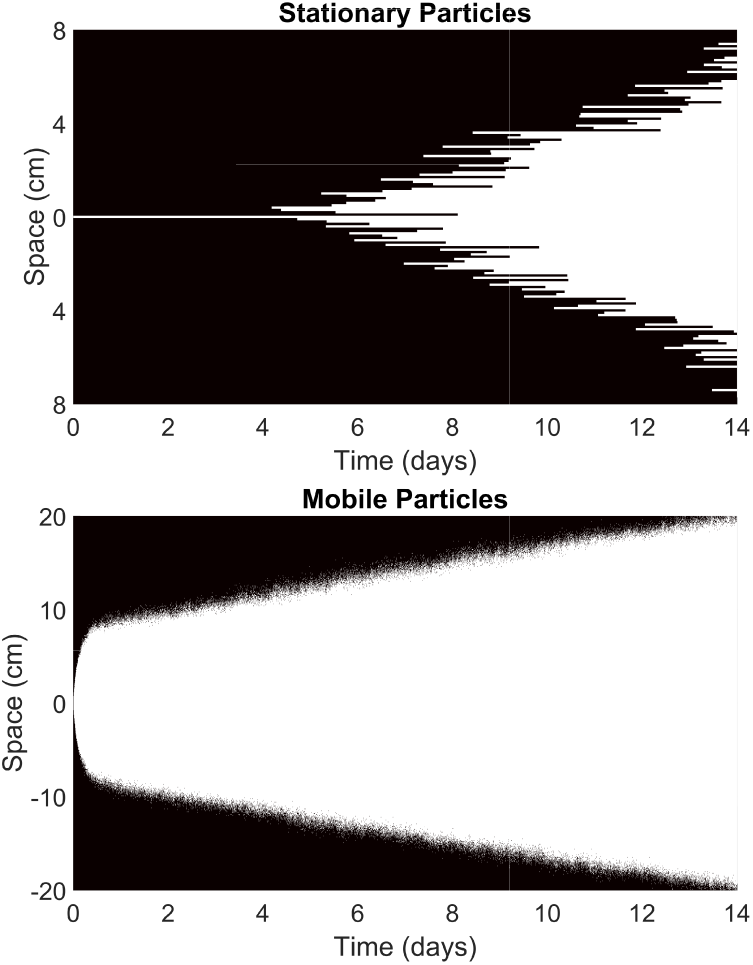
Manipulation of the plots from the left column of Figure 3 showing the areas that may be significantly affected by the primary tumor and its metastatic spread. More specifically, areas that have, on average, at least 0.025 particles over all 1000 simulations are colored white. Areas in which this is not the case are colored black. The top shows results for stationary particles, and the bottom for mobile particles.

Figure 2 shows a comparison between the exact dynamics from Theorem 3.2 (black curve) and the average dynamics over 1000 individual realizations of the stochastic model (blue curve). While the simulations all begin with a single stationary particle, the expected number of particles increases to approximately 50 within the first day. This rapid increase is due to the relatively high shedding rate, resulting in the rapid creation of mobile particles. The slow-down upon reaching 50 particles reflects the expected asymptotic ratio of moving to stationary particles (comment after Lemma 3.2) which is approximately 48 with the parameters from Table 2. With the metastatic reproduction number *R*_0_ ≈ 5 × 10^6^ ≫ 1, we expect the total particle number to grow exponentially at rate *α*_+_ ≈ 0.167 (Corollary 3.1). After an initial period of transience, we do see exponential growth, both in the exact and simulated results.

Figure 3 illustrates the average spatio-temporal dynamics of the stochastic model. The left column presents the full spatio-temporal dynamics of the stationary (top) and mobile (bottom) particles, while the right column shows the spatial distribution of the stationary (top) and mobile (bottom) particles at time *t* = 14 days. In no individual simulation did we see the original established tumor perish. This result is not unexpected given the probability of a tumor perishing over the 14 days considered in our simulations is 1 - exp(−14*δ*_2_) = 4.48 × 10^−7^. Consequently, we always have at least one stationary particle located at position *x* = 0. This explains both the horizontal line in the top left plot (note the difference in scales), as well as the tall bar at the origin in the top right plot. The histograms on the right side of Figure 3 show relatively symmetric distributions of both stationary and mobile particles centered around the origin. While the individual location of each particle is given by a normal distribution as a result of the Brownian dynamics, the distribution of the aggregate particles (both stationary and mobile) is not normal. The reason is that shedding, settlement and death cause additional randomness. Even when *λ* = 0 (no settlement) and *δ*_1_ = *δ*_2_ = 0 (no deaths) so that only one stationary particle sheds moving particles, the distribution of the aggregate moving particles will not be normal. This can be seen from (2.1), which becomes *F*_*t*_ = *μG*, hence the density of the aggregate moving particles in this case is an integral of normal densities and not a normal density itself.

In order to more clearly see the interface between empty space and invading cancer cells, we have taken the data in the left column of Figure 3, and simplified them to be either 1 if there was, on average, at least 0.025 particles in that location across all 1000 simulations, or 0 otherwise. The results of this simplification are presented in Figure 4. We can see that for stationary particles, it takes close to four days before we see any significant establishment events. This result mirrors the observations made by Cameron et al. (2000) that no metastases established in the first four days post-injection.

Following a rapid initial jump, the mobile particle boundary appears to invade at a more or less constant speed. These mobile ‘boundary’ dynamics (Figure 4) are in stark contrast with the ‘interior’ dynamics (bottom left in Figure 3, in particular, the blue-teal interface) where the level sets form triangular regions with edges whose slopes are increasing as we advance through time.

## 5. Conclusion and Discussion

We introduced and analyzed a *branching stochastic process with settlement* and we applied it to metastatic cancer growth. The fact that the expected number *F* of particles, the variance *V*, the distribution *H* of the furthest particle and the extinction probability *Q* satisfy the same type of integro-differential equation with distributed delay (2.5) reveals the recursive structure of this process. Methods from differential equations theory become available to analyze the qualitative behavior of this stochastic process. A recurring quantity was identified to play the role of a basic reproductive number, similar to epidemic models (Hethcote, 2000), which we call the *metastatic reproductive number*

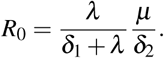

For the mouse data that we analyzed as an example, we found *R*_0_ ~ 10^6^, which, of course, is huge. This is expected, as cell lines for metastasis studies are chosen specifically to generate metastases efficiently and reliably.

The value of *R*_0_ for a typical human cancer will be quite different, and we leave a detailed estimate of *R*_0_ for human cancers for further studies. Still, we can already see the impact of various possible treatment strategies. To shrink *R*_0_, we like to reduce the shedding rate *μ* and the settlement rate *λ* while increasing the death rates *δ*_1_ and *δ*_2_ for moving and stationary particles, respectively. For example, the death rate *δ*_1_ for CTCs could be increased through platelet inhibitors. Platelets are known to shield cancer cells from the immune surveillance, and less platelets can make cancer cells more exposed and more vulnerable (Coupland et al., 2012; Riggi et al., 2018; Shahriyari, 2016). The settlement rate *λ* might be reduced through decreasing the availability of metastatic niches (Riggi et al., 2018). This can be achieved through very simple means such as reduced pH-levels of tissue (Silva et al., 2009) to very advanced means such as novel immunotherapies designed to disrupt the preparation of the premetastatic niche (Kaplan et al., 2005). However, removing 90% of cancer sites would not change the final outcome since the reproductive number is unchanged. A partial removal would significantly delay cancer spread, but metastasis would recur over time. Overall, the index *R*_0_ has the potential to become a useful quantity in treatment planning.

We see various extensions and limitations of the model as we discuss now.

1. While we started with a stationary individual, we could have started with a particle that is moving randomly, but this would make the analysis more difficult. Furthermore, if we know the function *F* of our model, we can find the corresponding function 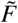 in a model with randomly moving first particle by

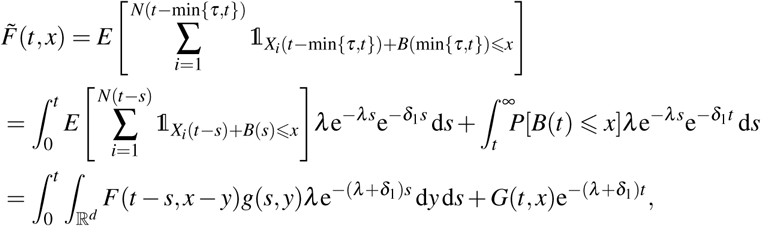

where *B*(min {*τ*,.}) is a random process describing the movement of the first particle up to time *τ*, and the factor 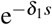 is the survival probability of the first particle at time *s*.
2. A classical related model is branching Brownian motion (Bovier, 2016), where all particles move according to Brownian motion and there are no deaths, no settlements and births occur independently at some exponentially distributed time *V* ~ Exp(*μ*). Similarly to the proof of Theorem 2.1, it can be shown that the expected location function 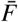 satisfies

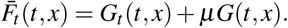 Note that this is different from our model even when assuming no deaths (*δ*_1_ = *δ*_2_ = 0) and no settlement (*λ* = 0). Indeed, for *δ*_1_ = *δ*_2_ = *λ* = 0, *F* in our model satisfies *F*_*t*_ (*t, x*) = *μG*(*t, x*). The reason for this difference to the case of branching Brownian motion is that, in our model with *δ*_1_ = *δ*_2_ = *λ* = 0, there is always a particle located at zero, and only this particle emits further particles.
3. In the one-dimensional, linear case with *r*_2_ = 0 and for constant decay rate *q*(*t*) = const., we can use Fourier and Laplace transforms to find an explicit solution of the general type (2.5) of non-local integro-differential equations. Let *ℱ, ℱ*^−1^ denote the Fourier transform and its inverse and let *ℒ*, *ℒ*^−1^ denote the Laplace transform and its inverse, respectively. We use the hat 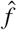 to denote the Fourier transform of a function and the tilde 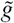 for the Laplace transform, i.e.

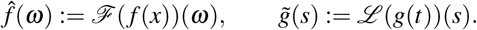 Transforming (2.5) in the case of *r*_2_ = 0 and *q*(*t*) =const. leads to

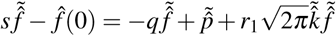

for an unknown function 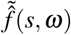 in Laplace-Fourier space. We can solve this algebraic equation and find

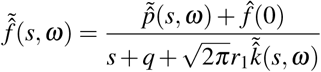 Hence the solution of (2.5) for *r*_2_ = 0 and *q*(*t*) = const. is:

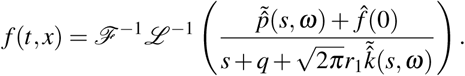 It has to be seen if this formula can shed valuable information about the branching stochastic process with settlement.
4. The spatial transport and settlement of a real cancer in a human body is much more complicated than assumed in our example. Here, as an example, we considered Brownian motion as spatial process and a homogeneous settlement rate, *λ*. However, our framework is based on a spatial process, (*B*(*t*))_*t⩾*0_, whose distribution could be more general and reflect more realistic body-wide properties. Such a specification is a complex issue and left for future research.
5. It is well known that certain tumors tend to metastasize to certain organs, for example prostate cancer preferentially metastasizes to the bone, and breast tumors often spread to the brain, bone, liver, and lungs (Chambers et al., 2002). In this case, the settlement rate, *λ*, is no longer homogeneous, rather it depends on the location *x*. Moreover, this spatial dependency encodes the locations of pre-metastatic niches (Kaplan et al., 2005). However, the branching stochastic process with settlement would then lose its recursive nature, which was crucial in the proofs of our results. Consequently, additional work must be completed before this intricacy can be included into future iterations of the model.

## Acknowledgements

We would like to thank two anonymous referees for valuable comments, which enabled us to significantly improve the paper. CF and TH gratefully acknowledge financial support by the Natural Sciences and Engineering Research Council of Canada through Discovery Grants. AR gratefully acknowledges funding from an Alberta Innovates Graduate Student Scholarship.

